# E484K mutation in SARS-CoV-2 RBD enhances binding affinity with hACE2 but reduces interactions with neutralizing antibodies and nanobodies: Binding free energy calculation studies

**DOI:** 10.1101/2021.02.17.431566

**Authors:** Wei Bu Wang, Yu Liang, Yu Qin Jin, Jing Zhang, Ji Guo Su, Qi Ming Li

## Abstract

The pandemic of the COVID-19 disease caused by SARS-CoV-2 has led to more than 100 million infections and over 2 million deaths worldwide. The progress in the developments of effective vaccines and neutralizing antibody therapeutics brings hopes to eliminate the threat of COVID-19. However, SARS-CoV-2 continues to mutate, and several new variants have been emerged. Among the various naturally-occurring mutations, the E484K mutation shared by both the 501Y.V2 and 501Y.V3 variants attracted serious concerns, which may potentially enhance the receptor binding affinity and reduce the immune response. In the present study, the molecular mechanism behind the impacts of E484K mutation on the binding affinity of the receptor-binding domain (RBD) with the receptor human angiotensin-converting enzyme 2 (hACE2) was investigated by using the molecular dynamics (MD) simulations combined with the molecular mechanics-generalized Born surface area (MMGBSA) method. Our results indicate that the E484K mutation results in more favorable electrostatic interactions compensating the burial of the charged and polar groups upon the binding of RBD with hACE2, which significantly improves the RBD-hACE2 binding affinity. Besides that, the E484K mutation also causes the conformational rearrangements of the loop region containing the mutant residue, which leads to more tight binding interface of RBD with hACE2 and formation of some new hydrogen bonds. The more tight binding interface and the new hydrogen bonds formation also contribute to the improved binding affinity of RBD to the receptor hACE2. In addition, six neutralizing antibodies and nanobodies complexed with RBD were selected to explore the effects of E484K mutation on the recognition of these antibodies to RBD. The simulation results show that the E484K mutation significantly reduces the binding affinities to RBD for most of the studied neutralizing antibodies, and the decrease in the binding affinities is mainly owing to the unfavorable electrostatic interactions caused by the mutation. Our studies revealed that the E484K mutation may improve the binding affinity between RBD and the receptor hACE2, implying more transmissibility of the E484K-containing variants, and weaken the binding affinities between RBD and the studied neutralizing antibodies, indicating reduced effectiveness of these antibodies. Our results provide valuable information for the effective vaccine development and antibody drugs design.

## INTRODUCTION

SARS-CoV-2 that causes the COVID-19 disease with high morbidity and mortality has spread rapidly across the world. According to the reports of the World Health Organization (WHO), SARS-CoV-2 has led to 93,194,922 infection cases and 2,014,729 deaths until 17 January 2021[1], which poses great threats to public health and brings heavy burdens to global economy. The significant progress in the developments of COVID-19 vaccines[2,3], including inactivated vaccine[4], mRNA vaccine[5,6], live vectorial vaccine[7,8] and recombinant protein subunit vaccine[9,10], as well as the neutralizing antibody therapeutics [11,12] is encouraging to prevent the pandemic of SARS-CoV-2. However, the virus is in constant evolution and several more contagious variants have emerged[13–16]. It is important to investigate the molecular mechanism for the impacts of the naturally occurring mutations on the infectivity of SARS-CoV-2, as well as on the immune effectiveness of the vaccines and the efficacy of the neutralizing antibodies.

It has been revealed that SARS-CoV-2 uses the spike (S) protein protruding on the surface of the virus membrane to bind to the receptor, i.e., human angiotensin-converting enzyme 2 (hACE2), on the host cells[17]. S is a homo-trimeric glycoprotein and each protomer is composed of S1 and S2 subunits. To recognize and bind with hACE2, the receptor-binding domain (RBD) of S1 undergoes a *“*down*”* to *“*up*”* conformational transition to expose the receptor-binding motif (RBM), which is directly involved in the interactions with the host cell receptor. Receptor binding triggers the dissociation of the S1 subunit and the transition of the S2 subunit from the prefusion to the postfusion states, which then leads to the membrane fusion and the invasion of the virus to the host cells[18]. Due to its critical function during the infection process, RBD predominantly determines the infectivity of SARS-CoV-2[19,20]. In addition, many studies have indicated that the S protein, especially the RBD, contains the major neutralizing epitopes, and the RBD-targeting antibodies immunodominantly contribute to the neutralizing activity of the convalescent sera from SARS-CoV-2 infected patients[21–23].

SARS-CoV-2 evolves continuously, and several new variants have already appeared and spread rapidly in many countries. On November 2020, a SARS-CoV-2 variant, termed as lineage B.1.1.7 or 501Y.V1, was detected in England, which rapidly became the dominant epidemic strain and spread to more than 30 countries[14,24,25]. Another new variant, named lineage B.1.351 or 501Y.V2, was discovered in South Africa on December 2020, which caused a new wave of SARS-CoV-2 infections and replaced other coronavirus strains to be one of the most pandemic[15]. In January 2021, a new variant named P.1 or 501Y.V3 was found in the travelers from Brazil, which has been widely prevalent in Amazonas state of north Brazil and has spread to Faroe Islands, South Korea and the United states[16,26]. Compared with the original SARS-CoV-2 strains, the 501Y.V1, 501Y.V2 and 501Y.V3 variants are potentially more transmissible, which may also reduce the effectiveness of the current vaccines and neutralizing antibodies[27]. The outbreaks of these new arising strains have attracted serious concerns globally. In these new epidemic strains, several residue mutations occur in the RBM of the S protein, which are directly involved in the interactions with the receptor hACE2 as well as with some immunodominant neutralizing antibodies. The 501Y.V1 strain carries the N501Y mutation in the RBM of the S protein. Experimental and simulation studies have revealed that the N501Y mutation improves the binding affinity of RBD to the receptor hACE2, which is believed to account for the more transmissibility of the 501Y.V1 variant than the original strains[28–32]. Fortunately, the sera neutralization assay showed that the N501Y mutation has little effect on the neutralization of the human sera elicited by the BNT162b2 mRNA vaccine[25]. In the 501Y.V2 and 501Y.V3 variants, besides the N501Y mutation, two other mutations E484K and K417N (or K417T) were occurred in RBD of the S protein, and it is revealed that the E484K mutation obviously enhances the binding affinity between RBD and hACE2[33].

More importantly, growing evidences indicated that the E484K mutation shared by both the 501Y.V2 and 501Y.V3 variants, named *“*escape mutation*”*, distinctly reduces the neutralization activity and even may escape from the neutralizing antibodies in the convalescent plasma of COVID-19 patients, which may weaken the effectiveness of the vaccines and the efficacy of the neutralizing antibody therapeutics in developments[21,31,34–36]. Understanding the molecular mechanism behind the effects of the E484K mutation on the RBD-hACE2 interactions as well as on the effectiveness of the neutralizing antibodies and nanobodies targeted at the wild-type RBD is of great significance for the developments of the therapeutic antibodies and vaccine designs.

In the present work, all-atom molecular dynamics (MD) simulation combined with the molecular mechanics-generalized Born surface area (MMGBSA) method[37,38] was employed to investigate the impacts of E484K mutation on the binding affinities of RBD with the receptor hACE2 as well as with the neutralizing antibodies and nanobodies. Our calculation results show that the E484K mutation may improve the binding affinity of RBD to the receptor hACE2 owing to more favorable electrostatic forces and more tight binding interface caused by the mutation, which implies more transmissibility of the E484K-containing variants. In addition, for most of the studied neutralizing antibodies and nanobodies, the E484K mutation leads to reduced binding affinities between RBD and these antibodies mainly due to the mutation-caused disadvantage electrostatic interactions, which weakens the effectiveness of these antibodies and even evade the immune protect.

## MATERIALS AND METHODS

### Protein structures preparation

The atomic coordinates file of the wild-type RBD complexed with the receptor hACE2 was downloaded from the Protein Data Bank (PDB) with the accession code 6M0J[17]. The structures of the complex formed by the wild-type RBD and various neutralizing antibodies and nanobodies were also obtained from PDB with accession codes listed in Table 1. To investigate the impacts of the E484K mutation on the binding affinity of RBD with hACE2 and the neutralizing antibodies, the residue Glu484 was replaced with Lys in these complex structures by using the UCSF chimera software to construct the corresponding mutant structures, in which the sidechain conformation of the mutated residue was chose as the rotamer with the highest probability according to the Dunbrack 2010 library[39].

**Table 1.**
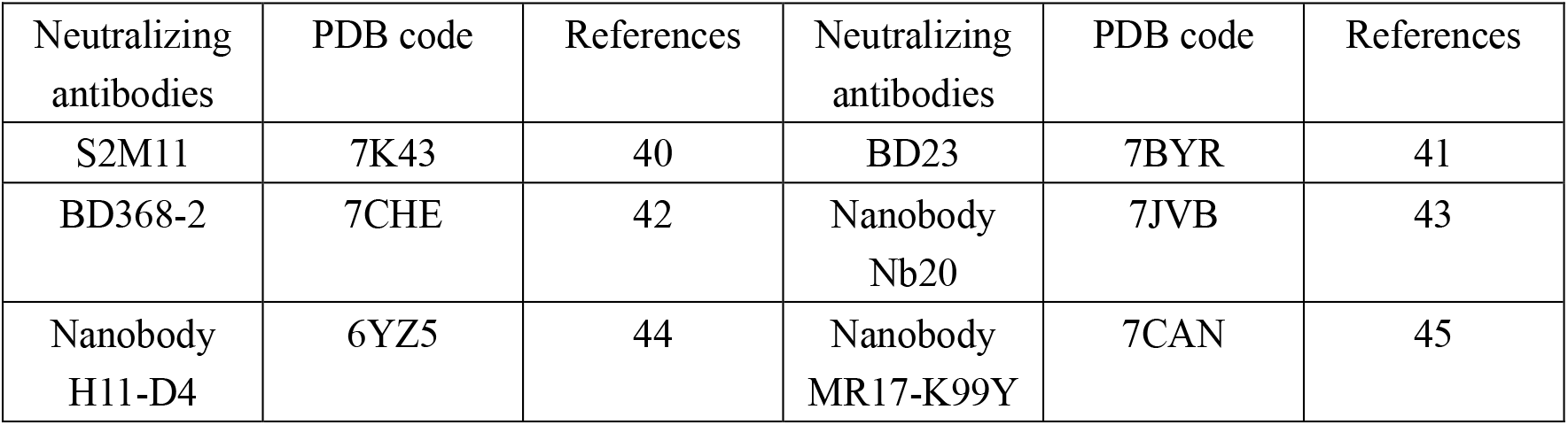
The complex formed by RBD and various neutralizing antibodies as well as nanobodies studied in this work

### Molecular dynamics (MD) simulations

The wild-type and the corresponding constructed mutant structures of RBD complexed with the receptor hACE2 as well as with the neutralizing antibodies were subjected to all-atomic MD simulations. In order to verify the convergence of our simulation and calculation results, a total of 5 independent MD simulations were carried out for each studied system. All the simulations were performed by using Amber16 software with the ff14SB fore field[46,47]. The input files for MD simulations were prepared with LEAP module of AmberTools16. After structure preparation, the simulation system was solvated in a cubic box with TIP3P water molecules and the size of the box was set to ensure that all protein atoms were at least 1.0 nm away from the box edges. The counter ions Na^+^ or Cl^-^ were added into the water box to neutralize the net charges of the system. Then, the energy of the system was minimized in two stages. In the first stage, the protein atoms were fixed by position restraints with a harmonic force constant of 10 kcal/(molÅ^2^), and only the water molecules as well as the counter-ions were energy minimized with 10000 steps of steepest descent algorithm followed by 10000 steps of conjugate gradient minimization algorithm. In the second stage, the entire system was energy minimized without any position restraints by using 10000 cycles of steepest descent algorithm combined with 10000 cycles of conjugate gradient minimization. After energy minimization, a 200ps constant volume (NTV ensemble) MD simulation with position restraints of 10 kcal/(molÅ^2^) on protein atoms was performed, in which the system was gradually heated to the temperature of 300 K. Subsequently, a 1 ns isothermal-isobaric (NTP ensemble) equilibration simulation at 300 K was carried out, in which the position restraints on protein structure were gradually released. In the equilibration simulation, the position restraints were gradually removed in five steps with the harmonic forces of 10.0, 5.0, 1.0, 0.5 and 0.25 kcal/(molÅ^2^), respectively, and each equilibration step was performed for 200 ps. Finally, the production NTP MD simulation without position restraints was performed for 6ns to obtain the simulation trajectory. For each studied system, the production NTP simulations were independently repeated five times. In these NTP MD simulations, the temperature of the system was kept at 300K with the Langevin coupling algorithm, and the pressure of the system was kept at 1 atm with Berendsen method. The cutoff value for the non-bonded van der Waals and short-range electrostatic interactions was set to 10 Å. All hydrogen bonds were constrained with the SHAKE algorithm and the integration time step for the simulations was set to 2 fs.

### Molecular mechanics-generalized Born surface area (MMGBSA) calculations

Based on the simulation trajectories, the binding free energy of RBD with the receptor hACE2 as well as with the various neutralizing antibodies was calculated by using the MMGBSA method. For each complex system, five binding free energy values were computed based on the five independent MD simulation trajectories to validate the convergence of our computation results. To investigate the impacts of the E484K mutation on the binding affinities of these complexes, the same MMGBSA calculations were carried out both for the systems formed by the wild-type RBD and the corresponding mutant systems formed by the RBD with the E484K mutation. According to the calculation results of the wild-type complexes and the corresponding mutants, the influences of the E484K mutation on the binding affinities of RBD with the receptor hACE2 as well as with the neutralizing antibodies were evaluated.

The MMGBSA calculations were carried out by using the Amber16 software with the single-trajectory method. In MMGBSA method, the binding free energy was estimated by summing the following contributions: Δ*G*_*bind*_ *=* Δ*E*_*MM*_ + Δ*G*_*polar*_ *+* Δ*G*_*nonPolar*_ − *T* Δ*S*, where Δ*E*_*MM*_ is the gas phase interaction energy including the electrostatic and the van der Waals interactions; Δ*G*_*Polar*_ is the polar contributions to the desolvation free energy, which is calculated with the GB equation; Δ*G*_*nonPolar*_ is the nonpolar component of the desolvation free energy, which is estimated empirically by using the solvent accessible surface area; *T*Δ*S* represents the change of the conformational entropy. In this study, the contribution of the conformational entropy was not considered, because the calculation of this term usually brings large errors for large-size systems like those studied in this work[48–52], which do not improve the calculation accuracy of the binding free energy. In order to explore the molecular mechanism for the influences of E484K mutation on the binding free energy, the MMGBSA calculation results were also decomposed into per-residue contributions.

## RESULTS AND DISCUSSION

### E484K mutation enhances the binding affinity between RBD and the receptor hACE2

The binding free energies of the wild-type RBD and the E484K mutant complexed with the receptor hACE2 were calculated, respectively, by using MD simulations combined with MMGBSA method. Then the difference in the binding free energy between the wild-type and the mutant systems was computed to investigate the impacts of the E484K mutation on the binding affinity of RBD with its receptor hACE2. In order to ensure the convergence of our calculation results, a total of five independent MD simulations as well as the corresponding MMGBSA calculations were performed both for the wild-type and the mutant complex systems. The calculation results are listed in Table 2. For most of these independent simulations, the binding free energies of the mutant RBD-hACE2 complex are distinctly lower than those of the wild-type. The average binding free energies for the wild-type and the mutant RBD-hACE2 complexes are −59.10 kcal/mol and −70.53 kcal/mol, respectively, which indicates that the E484K mutation significantly improves the binding affinity between RBD with its receptor hACE2. Our calculation results are consistent with the observed bioactivity changes, where the 501Y.V2 and 501Y.V3 SARS-CoV-2 strains with the E484K mutation display more transmissibility than the original strains. In MMGBSA method, the sum of the van der Waals and the nonpolar desolvation energies, i.e., Δ*E*_*vdw*_ *+* Δ*G*_*nonPolar*_, represents the hydrophobic interactions between the binding partners. The sum of the gas-phase electrostatic interactions and the polar desolvation energy, i.e., Δ*E*_*ele*_ *+* Δ*G*_*Polar*_, is responsible for the burial of the charged or polar groups upon binding. From Table 2, it is found that both for the wild-type and the mutant RBD-hACE2 complexes, the hydrophobic interactions are favorable, whereas the burial of the charged and polar groups is unfavorable. However, the E484K mutation significantly enhances the gas-phase electrostatic interactions and alleviates the unfavorable energies for the burial of the charged and polar groups upon the binding of RBD with hACE2 as shown in Table 2. The value of Δ*E*_*ele*_ is significantly decreased from −654.74 kcal/mol to −1113.71 kcal/mol and the Δ*E*_*ele*_ *+* Δ*G*_*Polar*_ energies is reduced from 44.54 kcal/mol to 35.99 kcal/mol, which mainly contributes to the improved binding affinity for the mutant system compared with the wild-type complex. Besides that, E484K mutation also leads to the decrease of the van der Waals and the nonpolar desolvation energies as shown in Table 2, where the value of Δ*E*_*vdw*_ *+* Δ*G*_*nonPolar*_ is reduce from −103.64 kcal/mol to −106.52 kcal/mol. Our results indicate that the mutation results in more tight binding between RBD and hACE2. The tighter binding induced by E484K mutation also contributes to the enhancements of the binding affinity for the mutant complex system.

**Table 2.** The binding free energies calculated by MMGBSA method for the wild-type RBD and the E484K mutant, respectively, complexed with the receptor hACE2 based on the five independent MD simulation trajectories^a)^.

| MD trajectories | $\Delta E_{vdw}$ | $\Delta E_{ele}$ | $\Delta G_{polar}$ | $\Delta G_{nonpolar}$ | $\Delta E_{vdw} + \Delta G_{nonpolar}$ | $\Delta E_{ele} + \Delta G_{polar}$ | $\Delta G_{bind}$ |
| --- | --- | --- | --- | --- | --- | --- | --- |
| WT<sup>b)</sup>-1 | -88.82 | -619.25 | 662.75 | -12.66 | -101.48 | 43.5 | -57.98 |
| WT-2 | -89.04 | -675.15 | 721.90 | -12.81 | -101.85 | 46.75 | -55.10 |
| WT-3 | -89.31 | -650.30 | 696.32 | -12.90 | -102.21 | 46.02 | -56.19 |
| WT-4 | -92.57 | -650.33 | 691.09 | -13.27 | -105.84 | 40.76 | -65.08 |
| WT-5 | -93.44 | -678.66 | 724.33 | -13.40 | -106.84 | 45.67 | -61.17 |
| WT-ave<sup>c)</sup> | -90.64 | -654.74 | 699.28 | -13.01 | -103.64 | 44.54 | -59.10 |
| MT<sup>d)</sup>-1 | -93.46 | -1124.49 | 1164.43 | -13.34 | -106.80 | 39.94 | -66.86 |
| MT-2 | -96.23 | -1134.52 | 1169.19 | -14.36 | -110.59 | 34.67 | -75.92 |
| MT-3 | -91.78 | -1111.58 | 1146.05 | -13.35 | -105.13 | 34.47 | -70.66 |
| MT-4 | -92.81 | -1105.55 | 1137.30 | -13.71 | -106.52 | 31.75 | -74.77 |
| MT-5 | -90.59 | -1092.43 | 1131.55 | -12.99 | -103.58 | 39.12 | -64.46 |
| MT-ave | -92.97 | -1113.71 | 1149.70 | -13.55 | -106.52 | 35.99 | -70.53 |

<sup>a)</sup>The unit of the values is kcal/mol; <sup>b)</sup>WT represents wild-type; <sup>c)</sup>ave denotes average; <sup>d)</sup>MT represents mutant.

By analysis of the binding interface around the mutated site of Glu484, it is found that there exist several charged residues on the receptor hACE2, including Glu35, Asp38 and Glu75 with negative charges, as well as Lys31 with positive charge (see Fig.1(a)). On average, the interface at the side of hACE2 exhibits electronegative character, which is unfavorable for the electrostatic interactions with the negatively charged Glu484. However, upon the replacement of Glu484 by Lys, these unfavorable gas-phase electrostatic interactions involving the residue 484 were converted to be favorable and thus the gas-phase electrostatic binding energy are significantly decreased as shown in Table 2, where the electrostatic energy is reduced from −654.74 kcal/mol to −1113.71 kcal/mol. In addition to the changes in the electrostatic interactions, we also investigate the mechanism for the decrease of the van der Waals and the polar desolvation energies caused by the E484K mutation. Based on the MD simulation trajectories, the average conformation in the MD simulation of the mutant complex was superposed onto that of the wild-type, as displayed in Fig. 1(b). Only the result for one of the five simulations was shown in Fig. 1(b), and the similar results were obtained for the other simulation trajectories (data not shown). It is found that the residue Glu484 is located on a flexible loop of RBD, and the substitution of Glu484 by Lys leads to obvious conformational movements of the flexible loop towards the receptor hACE2, which results in the more tight binding between RBD and hACE2. Our simulation results are consistent with the simulation results of Nelson[33]. Therefore, the conformational rearrangements of the binding interface caused by the mutation result in the decrease of the van der Waals and the polar desolvation energies. The conformational movements of the flexible loop may be due to the more favorable electrostatic forces between RBD and hACE2 in the mutant complex.

**Fig. 1.**
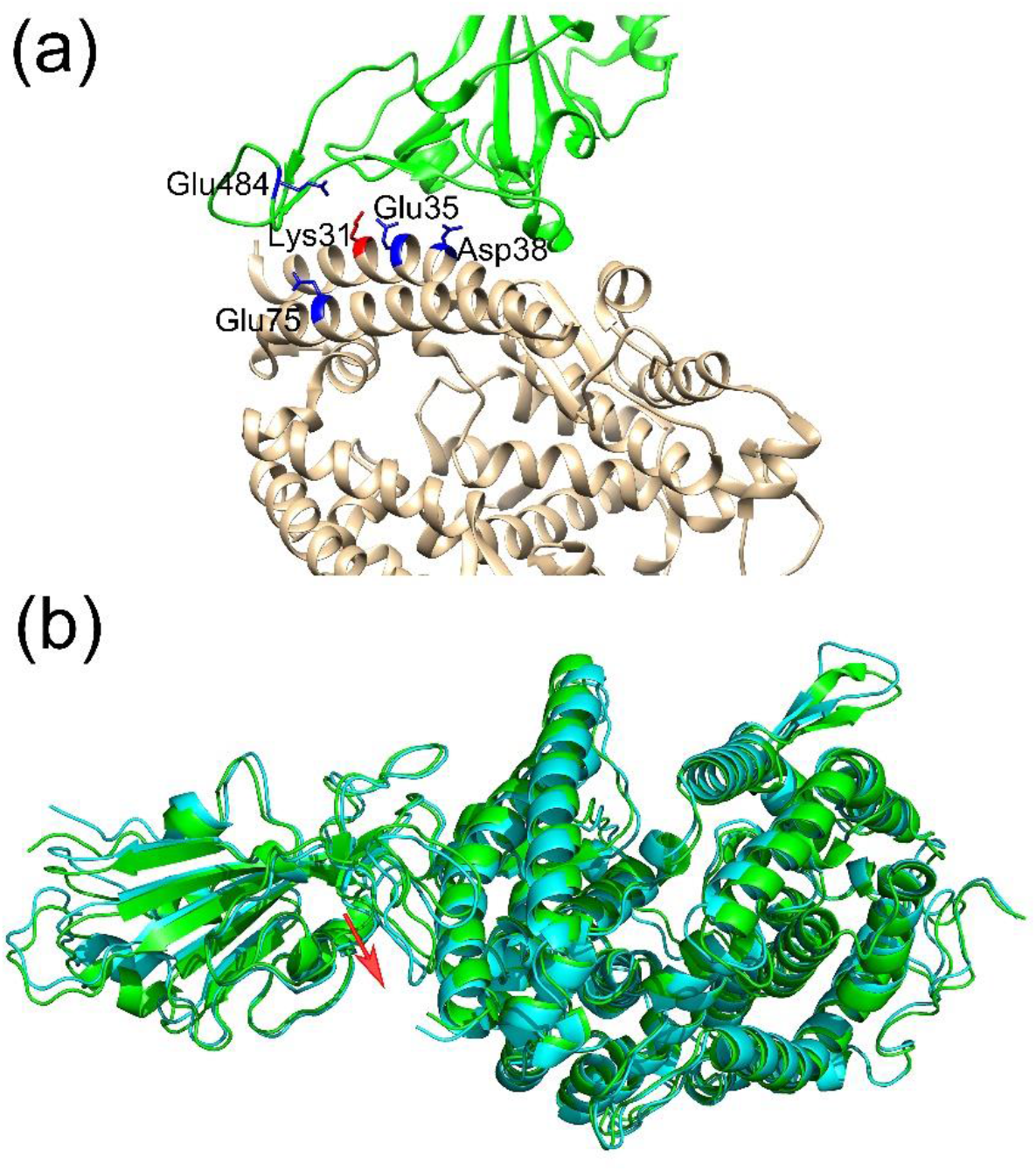
(a) The charged residues on hACE2 locating nearby the mutated residue E484K. In this figure, RBD and hACE2 are shown in green and brown colors, respectively. The positively and negatively charged residues are displayed with the stick model in red and blue colors, respectively. (b) The superposition of the average structure in the MD simulations between the wild-type complex and the E484K mutant. The average structures of the wild-type complex and the mutant are displayed in green and canyon colors, respectively. The red arrows in the figure represent the local structural movements of the mutant system compared with the wild-type.

In order to reveal the key residues responsible for the improvements of the binding affinity between RBD and hACE2, the per-residue energy decomposition analysis of MMGBSA was performed and the calculation results are displayed in Fig. 2. Only one of the five independent MD simulation trajectories was analyzed in the following studies, and the similar results can be obtained for the other simulation trajectories. Our calculation results show that in the wild-type RBD, the binding free energy contributed by the residue Glu484 is a positive value, which implies that the presence of Glu is unfavorable for the binding of RBD to hACE2 owing to the electrostatic repulsions as discussed above. Whereas, upon the substitution of Glu484 with Lys, the unfavorable electrostatic interactions were changed to be favorable and the binding free energy contributed by Glu484 became a negative value, as shown in Fig. 2. Besides the mutated residue E484K itself, the contributions to the binding free energy by the residues Tyr489, Gln493, Leu492, Phe490, Phe486 and Asn487 on RBD are also distinctly promoted, as shown in Fig. 2(a). At the side of hACE2, the residues Tyr83, Leu79 and Gln24 also exhibit improved contributions to the binding free energy, as displayed in Fig. 2(b). The improvements of the binding affinity contributed by these residues are mainly attributed to the more tight binding interface caused by the E484K mutation.

**Fig. 2.**
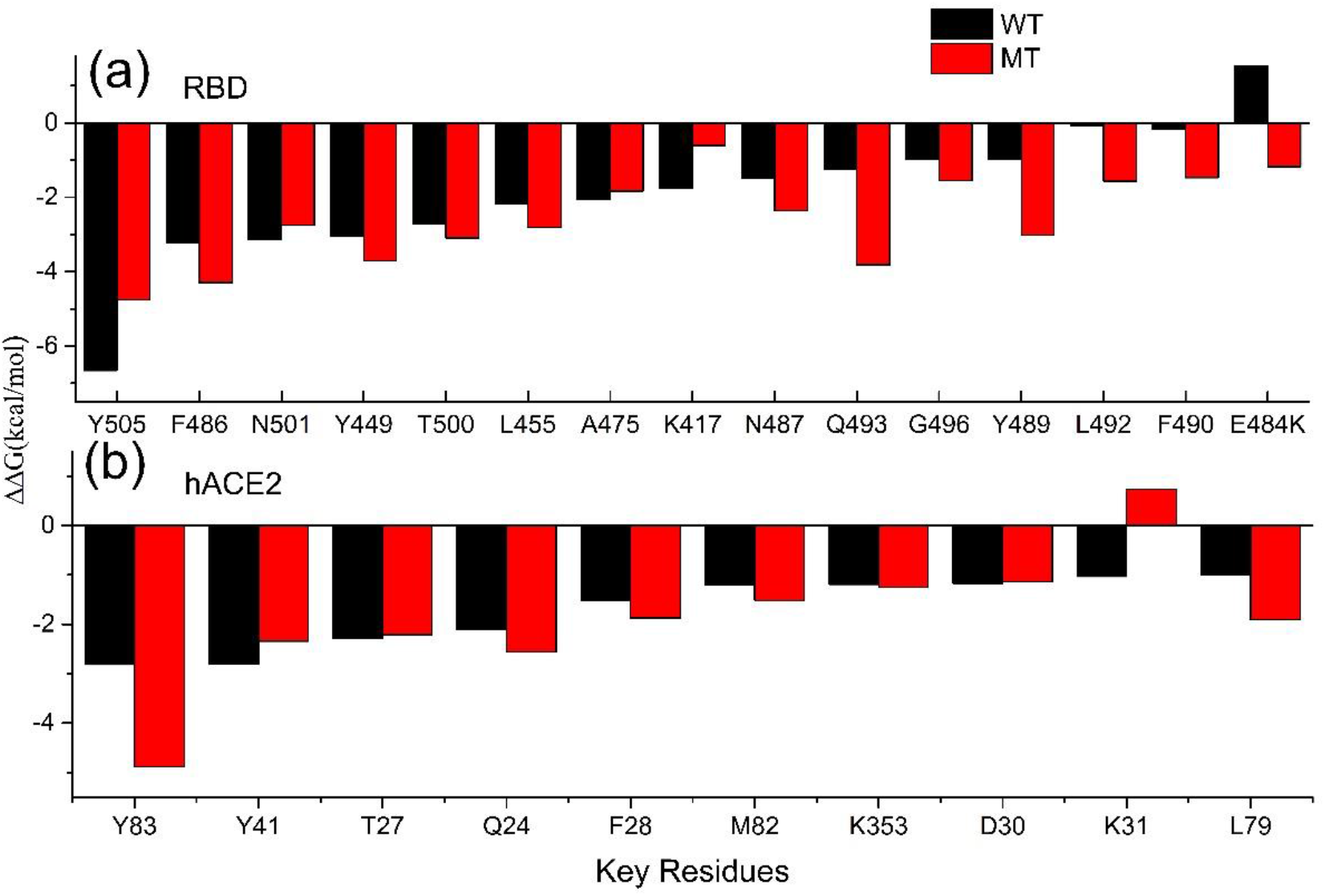
The binding free energies contributed by each of the key residues on RBD (a) and hACE2 (b), which are responsible for the improvements of the RBD-hACE2 binding affinity caused by the E484K mutation.

Based on the MD simulation trajectories, the conformational rearrangements around these residues were analyzed in detail. Our simulation results show that the conformational movements of the loop where the mutated residue is located also draw the region of residues 489-494 to move closer towards the receptor hACE2. Compared with the wild-type RBD-hACE2 complex structure, the sidechain of the residue Tyr489 moves closely towards the residue Tyr83 on hACE2 and form a new hydrogen bond between them in the mutant complex as shown in Table 3, in which the residues Tyr489 and Asn487 on RBD, and Gln24 and Tyr83 on hACE2 form a hydrogen bonds network in the mutant complex as shown in Fig. 3(a). The better packing of these residues and the new formed hydrogen bonds network between them account for the enhanced binding affinity of the mutant complex. The binding free energies of Tyr489 and Asn487 on RBD are decreased respectively from-0.97 kcal/mol and −1.48 kcal/mol to −3.02 kcal/mol and −2.35 kcal/mol, and those of Gln24 and Tyr83 on hACE2 are decreased from −2.11 kcal/mol and −2.81 kcal/mol to −2.56 kcal/mol and −4.88 kcal/mol, respectively. Fig. 3(b) shows that in the wild-type structure, the sidechains of the residues Leu492, Gln493 on RBD, and Lys31 on hACE2 are relatively far apart. Whereas, they move closer in the mutant structure and form new hydrogen bonds between them as shown in Table 3. The better packing of these residues and the more stable hydrogen bonds between them contribute to the stronger binding free energy for the residues Leu492 and Gln 493 in the mutant complex. Compared with the wild-type structure, the binding free energies of the residues Leu492 and Gln493 on RBD are changed from −0.088 kcal/mol and −1.25 kcal/mol to −1.57 kcal/mol and −3.82 kcal/mol, respectively. It should be noted that the E484K mutation may also cause decreased binding interactions for some residues. The electro-repulsive interactions between Lys31 on hACE2 and the mutated Lys484 lead to unfavorable contribution of Lys31 to the binding free energy in the mutant structure, as shown in Fig. 1(a) and Fig. 2. The decrease in the occupancy of the hydrogen bonds formed by Lys417 and Tyr505 with the corresponding residues in hACE2 results in the decreased contribution of these two residues to the binding affinity, as shown in Table 3 and Fig. 3(c). However, the unfavorable effects caused by the E484K mutation are minor. For most of the residues on the binding interface, the E484K mutation results in favorable electrostatic forces as well as more tight binding mode, which distinctly improves the binding affinity.

**Table 3.**
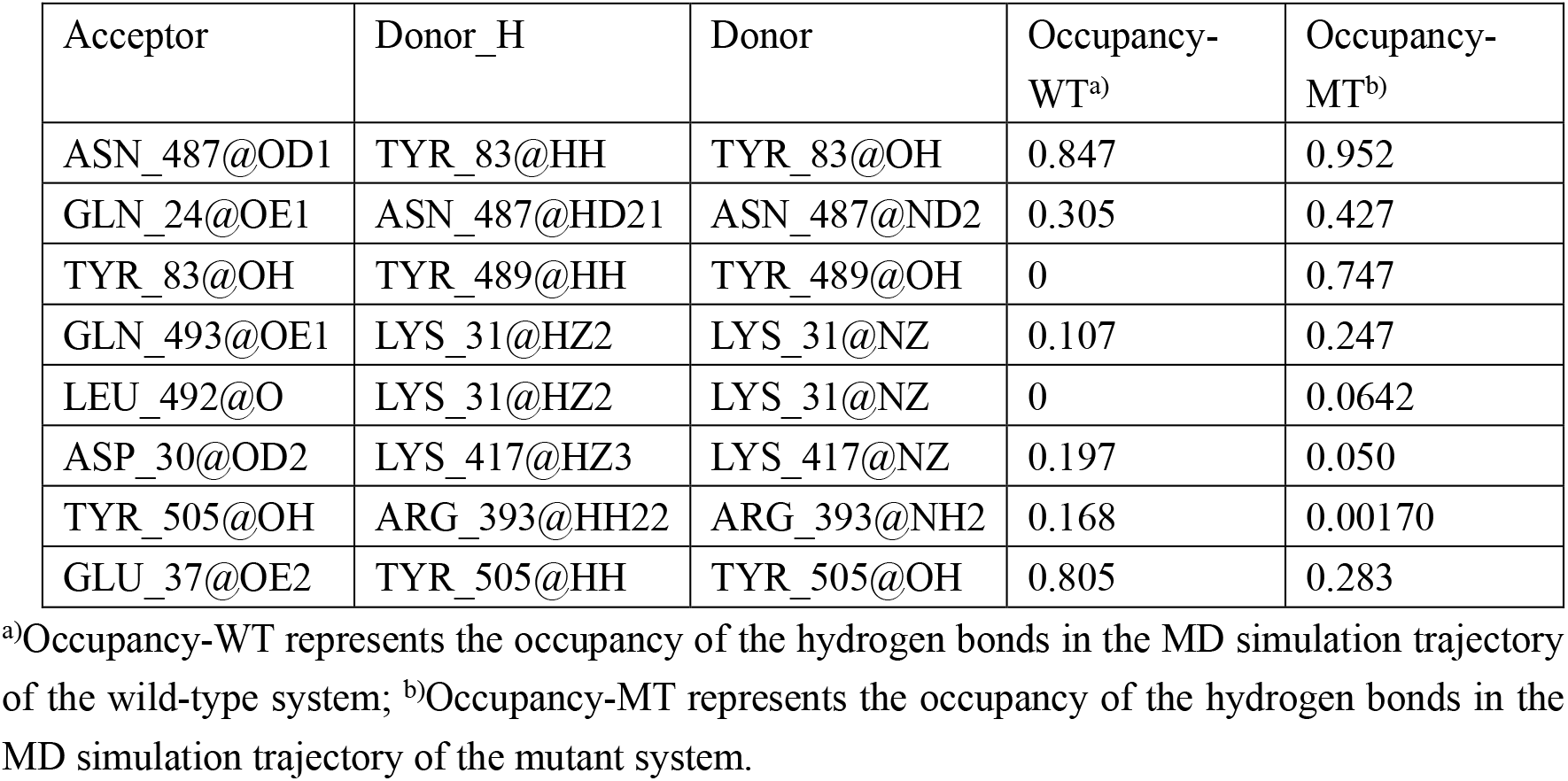
The E484K-induced changes in the occupancy of the hydrogen bonds formed by the residues discussed in the paper.

**Fig. 3.**
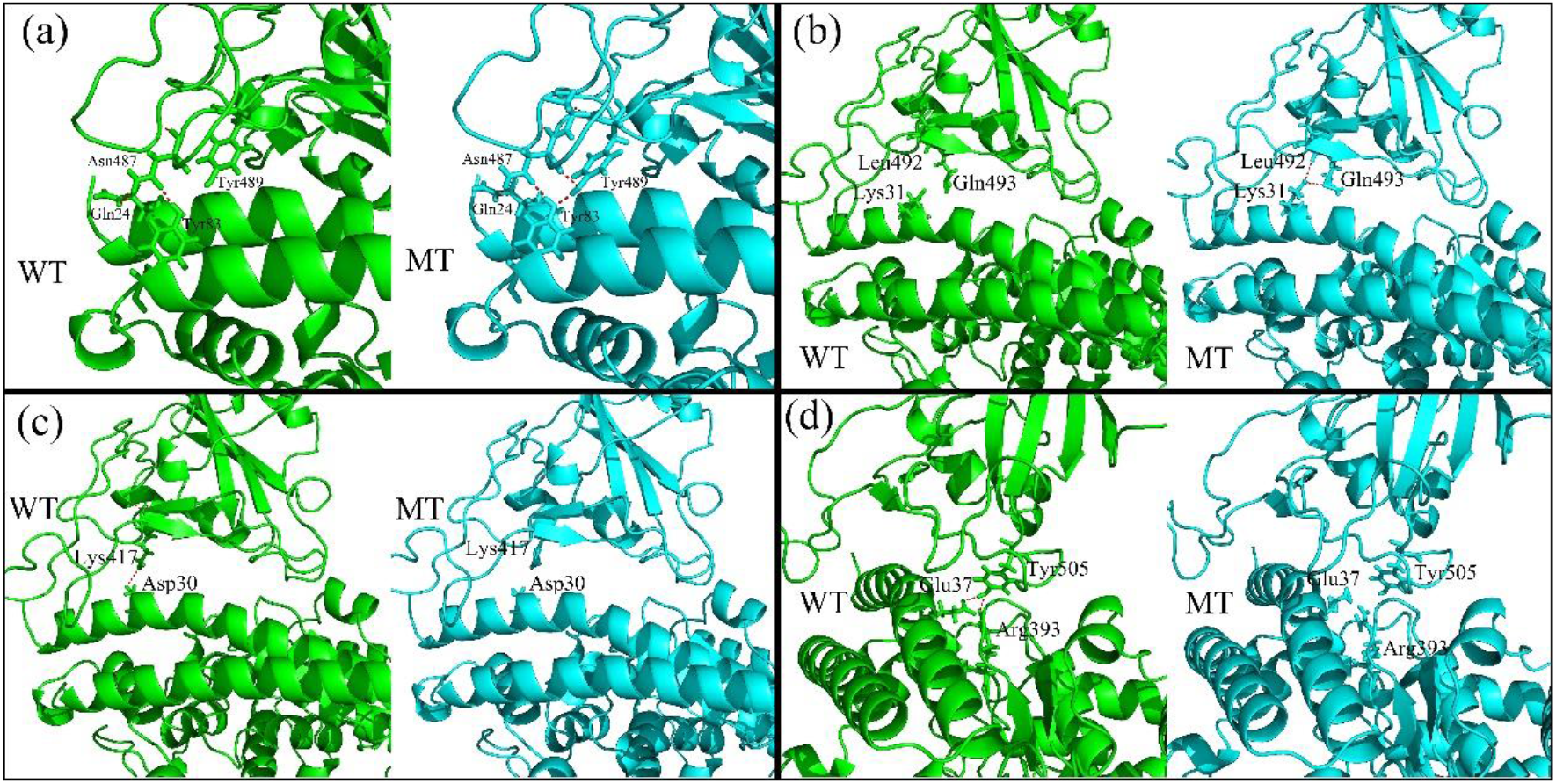
The local structural rearrangements of the RBD-hACE2 binding interface caused by the E484K mutation.

### E484K mutation distinctly reduces the binding affinity between RBD and most of the studied neutralizing antibodies and nanobodies

In the present study, a total of six neutralizing antibodies and nanobodies complexed with RBD were investigated to reveal the impacts of the E484K mutation on the binding affinities between them. For each RBD-antibody complex, five independent MD simulations were performed both for the wild-type and the mutant systems. Based on these simulation trajectories, the binding free energy of the wild-type RBD and the E484K mutant, respectively, complexed with these antibodies were calculated by using the MMGBSA method, and then the changes in the binding free energies between the wild-type and the mutant systems were computed to explore the influences of the E484K mutation on the binding affinities of RBD with these various neutralizing antibodies and nanobodies. The average binding free energy of the five MD simulations for each RBD-antibody complex is shown in Table 4, and the results for individual simulations are shown in Table S1. It is found that in these six studied neutralizing antibodies, five antibodies including S2M11, BD23, BD368-2, nanobody Nb20 and nanobody H11-D4, exhibit significantly decreased binding affinities with the E484K mutant compared with the wild-type RBD. The average binding free energies are increased from −58.02 kcal/mol, −54.24 kcal/mol, −48.41 kcal/mol, −64.06 kcal/mol and −47.77 kcal/mol to −41.52 kcal/mol, −32.96 kcal/mol, −39.53 kcal/mol, −48.79 kcal/mol and −37.15 kcal/mol for these antibodies, respectively. Our results indicate that for most of the studied neutralizing antibodies and nanobodies, the E484K mutation significantly reduces the binding affinities between these various antibodies and RBD, which may lessen the neutralization activity and even escape from these neutralizing antibodies. For nanobody MR17-K99Y, the E484K mutation has small effects on the binding affinity between this nanobody and RBD, where the binding free energy is changed from −70.71 kcal/mol to −71.22 kcal/mol. From the different components of the binding free energy as displayed in Table 4, it can be observed that for all these studied systems, the electrostatic energies are increased upon the mutation of E484K, which significantly penalize the binding of the antibodies to the mutant RBD. Especially, for the antibodies S2M11, BD23, BD368-2 and nanobody H11-D4, the electrostatic interaction is attractive in the wild-type complex where Glu with negative charge is present in the position at 484. Whereas, when Glu484 is mutated to Lys with positive charge, the attractive electrostatic interactions are changed to be repulsive as shown in Table 4, which are predominantly responsible for the decreased binding affinities of these antibodies with the E484K mutant RBD. For the other antibodies, the electrostatic energies were also raised in different degrees, which are disadvantaged for the binding of these antibodies with the mutant RBD. In summary, for most of the studied neutralizing antibodies, the E484K mutation significantly reduces the binding affinities between these antibodies with RBD, and the decrease of the binding affinities is mainly due to the repulsive electrostatic interactions caused by the mutation. Our results indicate that the E484K mutation may has disadvantageous effects on the neutralization activities and even may escape from the neutralization of these antibodies.

**Table 4.** The average binding free energies calculated by MMGBSA method for the wild-type RBD and the E484K mutant, respectively, complexed with various neutralizing antibodies and nanobodies based on the five MD simulation trajectories for each system^a)^.

| Neutralizing antibodies | RBD | $\Delta E_{vdw}$ | $\Delta E_{ele}$ | $\Delta G_{polar}$ | $\Delta G_{nonpolar}$ | $\Delta E_{vdw} + \Delta G_{nonpolar}$ | $\Delta E_{ele} + \Delta G_{polar}$ | $\Delta G_{bind}$ |
| --- | --- | --- | --- | --- | --- | --- | --- | --- |
| S2M11 | WT<sup>b)</sup> | -71.59 | -74.16 | 97.06 | -9.33 | -80.92 | 22.90 | -58.02 |
|  | MT<sup>c)</sup> | -72.79 | 31.18 | 8.59 | -8.50 | -81.29 | 39.77 | -41.52 |
| BD23 | WT | -71.28 | -122.61 | 149.54 | -9.89 | -81.17 | 26.93 | -54.24 |
|  | MT | -75.06 | 44.91 | 7.04 | -9.85 | -84.91 | 51.95 | -32.96 |
| BD368-2 | WT | -74.68 | -92.73 | 128.59 | -9.59 | -84.27 | 35.86 | -48.41 |
|  | MT | -70.23 | 3.98 | 35.93 | -9.21 | -79.44 | 39.91 | -39.53 |
| nanobody<br>Nb20 | WT | -82.75 | -237.52 | 268.30 | -12.09 | -94.84 | 30.78 | -64.06 |
|  | MT | -75.69 | -47.65 | 85.08 | -10.53 | -86.22 | 37.43 | -48.79 |
| Nanobody<br>H11-D4 | WT | -62.60 | -2.90 | 26.09 | -8.36 | -70.96 | 23.19 | -47.77 |
|  | MT | -64.60 | 271.64 | -236.30 | -7.89 | -72.49 | 35.34 | -37.15 |
| Nanobody<br>MR17-K99Y | WT | -102.33 | -299.26 | 344.51 | -13.63 | -115.96 | 45.25 | -70.71 |
|  | MT | -106.09 | -226.00 | 274.81 | -13.94 | -120.03 | 48.81 | -71.22 |

<sup>a)</sup>The unit of the values is kcal/mol; <sup>b)</sup>WT represents wild-type RBD; <sup>c)</sup>MT represents the mutant RBD.

In order to explore the physical mechanism and the related key residues responsible for the raising of the electrostatic energies caused by the E484K mutation, the changes of the binding free energy contributed by each residue as well as the mutation-induced local structural rearrangements were investigated for all these studied RBD-antibody systems based on the MD simulation trajectories and the per-residue energy decomposition analysis of the MMGBSA method. For each studied complex system, only one of the five independent MD simulation trajectories was analyzed in the following studies, and the similar results can be obtained for the other simulation trajectories.

For the neutralizing antibody BD23 complexed with RBD, the residue E484 of RBD forms attractive electrostatic interactions with the residue Arg107 on BD23 in the wild-type structure, as shown in Fig. 4(a). However, upon the mutation of GLu484 by Lys, the attractive electrostatic interactions with Arg107 are changed to be repulsive, which also leads to the sidechains of Lys484 and Arg107 moving apart. MMGBSA calculations show that the electrostatic energies contributed by the residues 484 and 107 are significantly increased from −78.55 kcal/mol and −4.18 kcal/mol to-1.62 kcal/mol and 31.28 kcal/mol (as displayed in Table 5), respectively, which are mainly responsible for the reduced binding affinity between the mutant RBD and the neutralizing antibody BD23.

**Table 5.** The binding free energy contributed by the key residues mainly responsible for the reduced binding affinity between RBD and the studied antibodies^a)^.

| Complex systems | Key residues | $\Delta E_{vdw}$ | $\Delta E_{ele}$ | $\Delta G_{polar}$ | $\Delta G_{nonpolar}$ | $\Delta E_{vdw} + \Delta G_{nonpolar}$ | $\Delta E_{ele} + \Delta G_{polar}$ | $\Delta G_{bind}$ |
| --- | --- | --- | --- | --- | --- | --- | --- | --- |
| S2M11-RBD-WT<sup>b)</sup> | Glu484 | -0.97 | -45.53 | 49.98 | -0.73 | -1.70 | 4.45 | 2.75 |
|  | Tyr33 | -0.40 | -9.37 | 5.70 | -0.15 | -0.55 | -3.67 | -4.22 |
|  | Asn52 | -0.78 | -10.65 | 7.32 | -0.08 | -0.86 | -3.33 | -4.19 |
|  | Ser55 | -0.78 | -10.16 | 5.25 | -0.34 | -1.12 | -4.91 | -6.03 |
| S2M11-RBD-MT<sup>c)</sup> | Lys484 | -3.33 | 0.33 | 2.93 | -0.68 | -4.01 | 3.26 | -0.75 |
|  | Tyr33 | -0.68 | -3.62 | 1.93 | -0.17 | -0.85 | -1.69 | -2.54 |
|  | Asn52 | -0.90 | -2.47 | 3.09 | -0.04 | -0.94 | 0.62 | -0.32 |
|  | Ser55 | -2.34 | -2.85 | 3.75 | -0.42 | -2.76 | 0.90 | -1.86 |
| BD23-RBD-WT | Glu484 | -0.85 | -78.55 | 81.16 | -0.70 | -1.55 | 2.61 | 1.06 |
|  | Arg107 | -1.24 | -4.18 | 5.19 | -0.20 | -1.44 | 1.01 | -0.43 |
| BD23-RBD-MT | Lys484 | -4.06 | -1.62 | 16.90 | -0.96 | -5.02 | 15.28 | 10.26 |
|  | Arg107 | -0.46 | 31.28 | -30.27 | -0.04 | -0.5 | 1.01 | 0.51 |
| BD368-2-RBD-WT | Glu484 | -2.77 | -78.78 | 78.66 | -0.67 | -3.44 | -0.12 | -3.56 |
|  | Arg100 | -0.97 | -12.06 | 12.17 | -0.31 | -1.28 | 0.11 | -1.17 |
|  | Arg102 | -4.74 | -27.37 | 22.76 | -0.76 | -5.50 | -4.61 | -10.11 |
|  | Asp106 | -1.70 | 1.84 | 4.15 | -0.08 | -1.78 | 5.99 | 4.21 |
| BD368-2-RBD-MT | Lys484 | -2.23 | -5.11 | 5.81 | -0.73 | -2.96 | 0.70 | -2.26 |
|  | Arg100 | -0.85 | 32.88 | -31.60 | -0.28 | -1.13 | 1.28 | 0.15 |
|  | Arg102 | -3.58 | 39.19 | -38.68 | -0.59 | -4.17 | 0.51 | -3.66 |
|  | Asp106 | -2.14 | -57.15 | 62.42 | -0.24 | -2.38 | 5.27 | 2.89 |
| nanobody Nb20-RBD-WT | Glu484 | -3.12 | -91.86 | 95.06 | -0.80 | -3.92 | 3.20 | -0.72 |
|  | Arg31 | -2.24 | -20.02 | 16.62 | -0.31 | -2.55 | -3.40 | -5.95 |
|  | Arg97 | -2.44 | -24.04 | 21.48 | -0.57 | -3.01 | -2.56 | -5.57 |
| nanobody Nb20-RBD-MT | Lys484 | -3.64 | 1.27 | 2.93 | -0.93 | -4.57 | 4.20 | -0.37 |
|  | Arg31 | -1.05 | 35.47 | -34.34 | -0.12 | -1.17 | 1.13 | -0.04 |
|  | Arg97 | -2.09 | 30.25 | -30.40 | -0.52 | -2.61 | -0.15 | -2.76 |
| Nanobody H11-D4-RBD-WT | Glu484 | -1.44 | -90.08 | 95.49 | -0.66 | -2.10 | 5.41 | 3.31 |
|  | Arg52 | -1.49 | -23.82 | 16.93 | -0.28 | -1.77 | -6.89 | -8.66 |
| Nanobody H11-D4-RBD-MT | Lys484 | -1.84 | 50.26 | -47.71 | -0.48 | -2.32 | 2.55 | 0.23 |
|  | Arg52 | -1.81 | 42.34 | -40.46 | -0.23 | -2.04 | 1.88 | -0.16 |
| Nanobody MR17-K99Y-RBD-WT | Glu484 | -3.09 | -24.95 | 26.94 | -0.39 | -3.48 | 1.99 | -1.49 |
|  | Arg33 | -3.38 | -3.33 | -1.26 | -0.55 | -3.93 | -4.59 | -8.52 |
|  | Arg59 | -6.22 | -16.40 | 15.68 | -0.78 | -7.00 | -0.72 | -7.72 |
|  | Lys65 | -2.23 | -12.57 | 15.83 | -0.54 | -2.77 | 3.26 | 0.49 |
| Nanobody MR17-K99Y-RBD-MT | Lys484 | -3.93 | 7.71 | -4.48 | -0.73 | -4.66 | 3.23 | -1.43 |
|  | Arg33 | -3.14 | 21.69 | -26.01 | -0.51 | -3.65 | -4.32 | -7.97 |
|  | Arg59 | -6.63 | 35.05 | -33.08 | -0.81 | -7.44 | 1.97 | -5.47 |
|  | Lys65 | -1.16 | 31.51 | -28.78 | -0.36 | -1.52 | 2.73 | 1.21 |

<sup>a)</sup>The unit of the values is kcal/mol; <sup>b)</sup>WT represents wild-type; <sup>c)</sup>MT represents mutant.

**Fig. 4.**
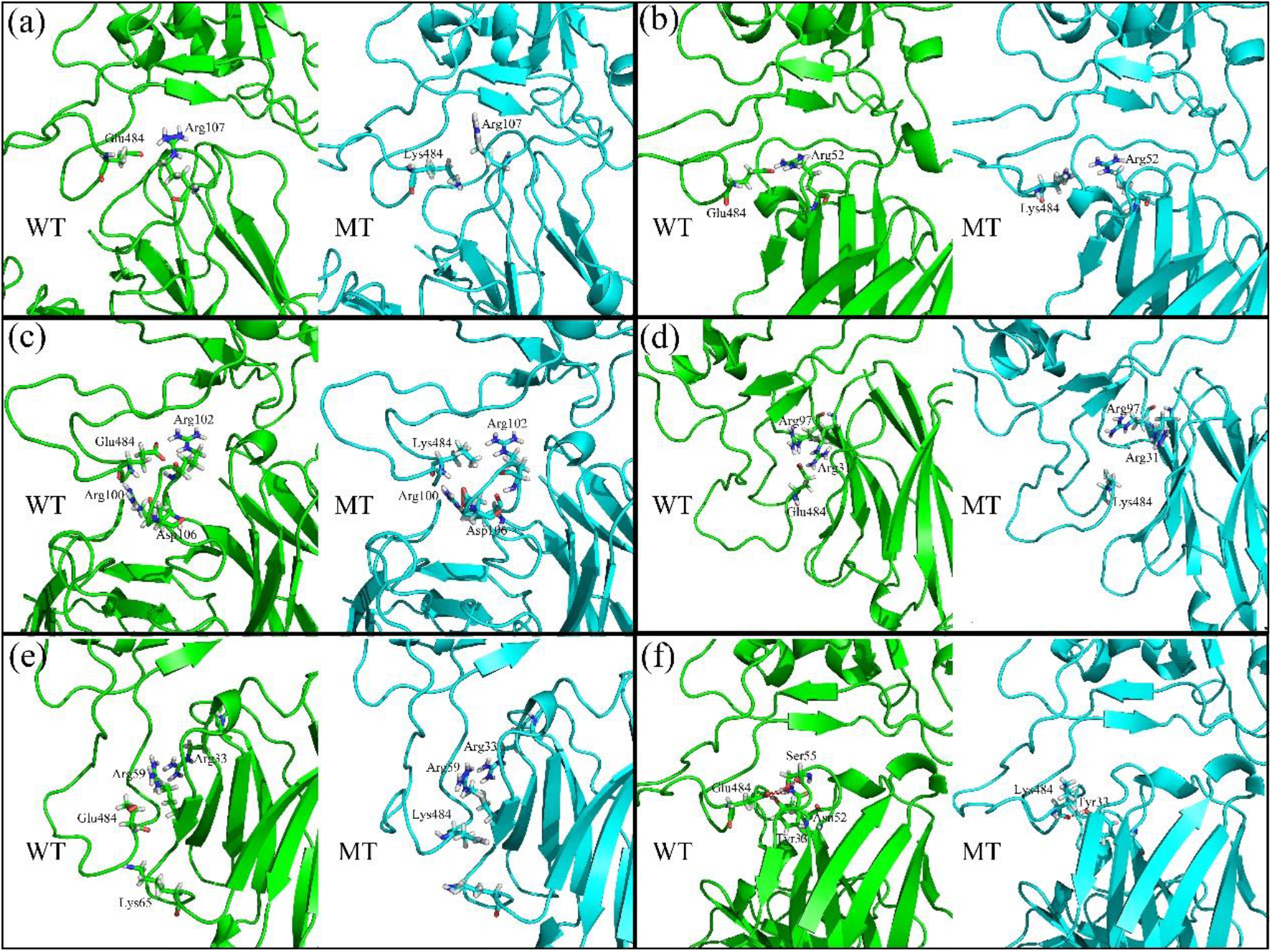
The locations as well as the conformational rearrangements of the key residues responsible for the raising of the electrostatic energies caused by the E484K mutation for the complex systems formed by RBD with the neutralizing antibody BD23 (a), the nanobody H11-D4 (b), the neutralizing antibody BD368-2(c), the nanobody Nb20 (d), the nanobody MR17-K99Y (e) and the neutralizing antibody S2M11 (f), respectively.

A similar situation is also observed for the nanobody H11-D4, where the positively charged residue Arg52 provides attractive electrostatic interactions with Glu484 of RBD in the wild-type complex structure as shown in Fig. 4(b). But the attractive electrostatic interactions are reversed to be repulsive when the negatively charged residue Glu484 is replaced by the positively charged Lys, which pushes these two residues away and weakens the binding affinity between RBD and the nanobody H11-D4.

For the neutralizing antibody BD368-2 complexed with RBD, the negatively charged residue Glu484 of RBD is sandwiched between two positively charged residues Arg100 and Arg102 of the antibody BD368-2, and the electrostatic interactions between them contribute to the stability of this kind of local structural arrangements, as shown in Fig. 4(c). Whereas, when the negatively charged residue Glu484 is substituted by Lys with positive charges, the electrostatic repulsions exerted by Arg100 and Arg102 lead to the swing away of the sidechain of Lys484. The electrostatic energies contributed by these charged residues are significantly increased in the mutant complex compared with the wild-type system as shown in Table 5, which mainly account for the reduced binding affinity of the antibody BD368-2 with the mutant RBD. There also exists a negatively charged residue Asp106 in BD368-2, which is more preferred for the binding of the mutant RBD compared to the wild-type. But the impact of Asp106 is relatively small owing to its large distance to the mutant residue E484K.

In the complex of nanobody Nb20 with RBD, upon the mutation of E484 by Lys, the attractive electrostatic interactions formed by Glu484 on RBD with Arg31 and Aeg97 on Nb20 are changed to be repulsive, which leads to the sidechain of Lys484 swing to the opposite direction, as shown in Fig. 4(d). The repulsive electrostatic interactions combined with the local structural rearrangements contribute to the reduced binding affinity between Nb20 and RBD, as indicated in Table 5. For all the above neutralizing antibodies, a similar physical mechanism was observed to be responsible for the reduced RBD-antibody binding affinity caused by the mutation E484K, where the attractive electrostatic interactions are converted to be repulsive resulted by the residue mutation with reverse charges.

For nanobody MR17-K99Y complexed with RBD, the positively charged residues Arg59, Arg33 and Lys65 on MR17-K99Y are electrostatically attractive with Glu484 on RBD in the wild-type complex, as shown in Fig. 4(e). The E484K mutation results in the convertion of these electrostatic interactions from attraction to repulsion, which also leads to the sidechain of Lys484 swing towards the concave of the antibody, as displayed in the right sub-figure in Fig. 4(e). The sidechain swing of Lys484 results in better packing of the binding interface and thus decreases the van der Waals energy, which compensates the increase of the electrostatic energy as indicated in Table 4 and Table 5. In total, the E484K mutation has little effect on the binding affinity between the nanobody MR17-K99Y and RBD due to the compensation of the impacts on the electrostatic interactions and van der Waals interactions.

For the complex structure of the neutralizing antibody S2M11 with RBD, the residue Glu484 forms several hydrogen bonds with the polar residues Ser55, Asn52 and Tyr33 of S2M11 in the wild-type system. When the residue Glu484 is substituted by Lys in the mutant system, most of these hydrogen bonds are destroyed and the electrostatic energies are significantly increased, as shown in Fig. 4(f) and Table 5. Although the van der Waals energies are decreased, the decrease of the van der Waals energies cannot compensate the increase of the electrostatic energies. Taken together, the E484K mutation obviously reduces the binding affinity between the neutralizing antibody S2M11 and RBD.

## CONCLUSION

SARS-CoV-2 evolves continuously, and several new variants have emerged and widely spread worldwide. It raised the worries about whether the new occurred variants would exhibit increased transmissibility or evade the immune protect. To date, most of the naturally occurring mutations have limited effects on the transmissibility, virulence and immune evasion of SARS-CoV-2. However, serious attentions have been paid to the E484K mutation shared both by the 501Y.V2 and 501Y.V3 variants initially discovered respectively in the United Kingdom and Brazil, which is considered to be potentially associated with improved transmissibility as well as decreased neutralization activity of the present antibodies. Understanding the molecular mechanism behind the impacts of the E484K mutation on the binding affinities of RBD with the receptor hACE2 as well as with various neutralizing antibodies is of great significance for the developments of effective vaccines and antibody drugs.

In the present work, the MD simulations combined with MMGBSA method were employed to evaluate the receptor binding free energy both for the wild-type RBD and the E484K mutant, and then the effects of the mutation on the binding affinity of RBD with the receptor hACE2 were investigated. The calculation results show that the upon the substitution of Glu484 with Lys, the unfavorable gas-phase electrostatic interactions between E484 and the negatively charged residues in hACE2 were converted to be favorable, which significantly contributes to the improved binding affinity of RBD with hACE2. Besides that, the E484K mutation also causes conformational rearrangements of the local structure around the mutant residue, which results in more tight binding interface of RBD with hACE2 and formation of some new hydrogen bonds. The better packing of the binding interface and the formation of new hydrogen bonds also lead to the enhancements of the binding affinity for the mutant complex system. The tighter RBD-receptor binding affinity caused by the mutation may be responsible for the more transmissibility of the E484K-containing variants. Additionally, in order to explore the impacts of the E484K mutation on the recognition of neutralizing antibodies to RBD, six different neutralizing antibodies and nanobodies complexed with RBD were studied, in which the changes of the binding free energy induced by the mutation were calculated by using MD simulations with MMGBSA method. The calculation results revealed that for most of the neutralizing antibodies and nanobodies, the E484K mutation significantly reduces the binding affinities between RBD and these antibodies, which may weaken the effectiveness of these antibodies and even evade the immune protect. The decrease in the binding affinities is mainly resulted by the mutation-caused unfavorable electrostatic interactions between RBD and the studied neutralizing antibodies. Our studies provide valuable information for the developments of effective vaccines and therapeutic antibodies.

