## Supplemental Table S1 for "E484K mutation in SARS-CoV-2 RBD enhances binding affinity with hACE2 but reduces interactions with neutralizing antibodies and nanobodies: Binding free energy calculation studies"

**Supporting Information**

Table S1. The calculated binding free energies of the five MD simulations both for the wild-type RBD and the E484K mutant, respectively, complexed with various neutralizing antibodies and nanobodies^a)^.

| Neutralizing antibodies | RBD | $\Delta E_{vdw}$ | $\Delta E_{ele}$ | $\Delta G_{polar}$ | $\Delta G_{nonpolar}$ | $\Delta E_{vdw}+\Delta G_{nonpolar}$ | $\Delta E_{ele}+\Delta G_{polar}$ | $\Delta G_{bind}$ |
| --- | --- | --- | --- | --- | --- | --- | --- | --- |
| S2M11 | WT^b)^-1 | -72.29 | -71.41 | 98.34 | -9.50 | -81.79 | 26.93 | -54.86 |
|  | WT-2 | -71.07 | -72.95 | 94.55 | -9.54 | -80.61 | 21.60 | -59.01 |
|  | WT-3 | -71.07 | -85.19 | 104.31 | -9.23 | -80.30 | 19.12 | -61.18 |
|  | WT-4 | -71.69 | -66.35 | 91.62 | -9.15 | -80.84 | 25.27 | -55.57 |
|  | WT-5 | -71.84 | -74.90 | 96.47 | -9.23 | -81.07 | 21.57 | -59.50 |
|  | MT^c)^-1 | -68.94 | 39.78 | -4.97 | -8.01 | -76.95 | 34.81 | -42.14 |
|  | MT-2 | -76.81 | 18.20 | 21.16 | -8.84 | -85.65 | 39.36 | -46.29 |
|  | MT-3 | -76.59 | 29.57 | 13.49 | -9.20 | -85.79 | 43.06 | -42.73 |
|  | MT-4 | -76.24 | 31.80 | 11.58 | -8.79 | -85.03 | 43.38 | -41.65 |
|  | MT-5 | -65.36 | 36.52 | 1.70 | -7.68 | -73.04 | 38.22 | -34.82 |
| BD23 | WT-1 | -66.64 | -94.78 | 122.53 | -9.36 | -76.00 | 27.75 | -48.25 |
|  | WT-2 | -65.19 | -112.90 | 137.60 | -9.05 | -74.24 | 24.70 | -49.54 |
|  | WT-3 | -69.59 | -142.07 | 164.08 | -9.85 | -79.44 | 22.01 | -57.43 |
|  | WT-4 | -80.14 | -113.78 | 148.51 | -10.57 | -90.71 | 34.73 | -55.98 |
|  | WT-5 | -74.86 | -149.52 | 175.00 | -10.64 | -85.50 | 25.48 | -60.02 |
|  | MT-1 | -75.20 | 48.37 | 3.46 | -9.75 | -84.95 | 51.83 | -33.12 |
|  | MT-2 | -73.27 | 30.91 | 25.35 | -9.33 | -82.60 | 56.26 | -26.34 |
|  | MT-3 | -88.62 | 14.76 | 46.24 | -11.91 | -100.53 | 61.00 | -39.53 |
|  | MT-4 | -67.57 | 68.74 | -21.35 | -9.05 | -76.62 | 47.39 | -29.23 |
|  | MT-5 | -70.66 | 61.79 | -18.50 | -9.18 | -79.84 | 43.29 | -36.55 |
| [BD368-2](http://www1.rcsb.org/structure/7CHH) | WT-1 | -73.09 | -93.82 | 131.00 | -9.53 | -82.62 | 37.18 | -45.44 |
|  | WT-2 | -75.32 | -99.93 | 137.89 | -9.63 | -84.95 | 37.96 | -46.99 |
|  | WT-3 | -71.12 | -69.56 | 105.08 | -9.03 | -80.15 | 35.52 | -44.63 |
|  | WT-4 | -76.48 | -85.18 | 122.90 | -9.76 | -86.24 | 37.72 | -48.52 |
|  | WT-5 | -77.37 | -115.18 | 146.11 | -9.99 | -87.36 | 30.93 | -56.43 |
|  | MT-1 | -71.69 | -10.90 | 49.65 | -9.34 | -81.03 | 38.75 | -42.28 |
|  | MT-2 | -74.12 | 12.88 | 30.87 | -9.81 | -83.93 | 43.75 | -40.18 |
|  | MT-3 | -71.76 | -3.45 | 41.75 | -9.40 | -81.16 | 38.3 | -42.86 |
|  | MT-4 | -61.54 | 12.48 | 23.97 | -8.18 | -69.72 | 36.45 | -33.27 |
|  | MT-5 | -72.04 | 8.89 | 33.40 | -9.30 | -81.34 | 42.29 | -39.05 |
| nanobody Nb20 | WT-1 | -82.47 | -215.50 | 250.00 | -11.74 | -94.21 | 34.50 | -59.71 |
|  | WT-2 | -79.09 | -255.69 | 276.06 | -11.95 | -91.04 | 20.37 | -70.67 |
|  | WT-3 | -87.38 | -222.60 | 260.15 | -12.81 | -100.19 | 37.55 | -62.64 |
|  | WT-4 | -82.20 | -225.04 | 260.52 | -11.84 | -94.04 | 35.48 | -58.56 |
|  | WT-5 | -82.60 | -268.79 | 294.75 | -12.11 | -94.71 | 25.96 | -68.75 |
|  | MT-1 | -70.11 | -65.04 | 99.20 | -10.15 | -80.26 | 34.16 | -46.10 |
|  | MT-2 | -77.66 | 12.32 | 28.76 | -10.10 | -87.76 | 41.08 | -46.68 |
|  | MT-3 | -71.57 | -67.14 | 100.65 | -10.29 | -81.86 | 33.51 | -48.35 |
|  | MT-4 | -87.44 | -53.01 | 94.14 | -11.78 | -99.22 | 41.13 | -58.09 |
|  | MT-5 | -71.68 | -65.37 | 102.64 | -10.36 | -82.04 | 37.27 | -44.77 |
| Nanobody H11-D4 | WT-1 | -61.10 | -3.53 | 23.71 | -8.30 | -69.40 | 20.18 | -49.22 |
|  | WT-2 | -66.17 | -13.59 | 41.41 | -8.57 | -74.74 | 27.82 | -46.92 |
|  | WT-3 | -62.76 | -1.09 | 24.00 | -8.28 | -71.04 | 22.91 | -48.13 |
|  | WT-4 | -60.30 | 3.45 | 18.18 | -8.05 | -68.35 | 21.63 | -46.72 |
|  | WT-5 | -62.65 | 0.26 | 23.15 | -8.57 | -71.22 | 23.41 | -47.81 |
|  | MT-1 | -63.51 | 270.39 | -237.76 | -7.75 | -71.26 | 32.63 | -38.63 |
|  | MT-2 | -65.73 | 289.51 | -252.18 | -8.05 | -73.78 | 37.33 | -36.45 |
|  | MT-3 | -64.75 | 267.61 | -232.46 | -7.95 | -72.70 | 35.15 | -37.55 |
|  | MT-4 | -66.63 | 258.85 | -222.20 | -8.13 | -74.76 | 36.65 | -38.11 |
|  | MT-5 | -62.37 | 271.84 | -236.88 | -7.56 | -69.93 | 34.96 | -34.97 |
| Nanobody MR17-K99Y | WT-1 | -103.48 | -328.47 | 366.24 | -13.86 | -117.34 | 37.77 | -79.57 |
|  | WT-2 | -107.75 | -280.44 | 330.06 | -14.31 | -122.06 | 49.62 | -72.44 |
|  | WT-3 | -107.07 | -329.92 | 372.20 | -14.34 | -121.41 | 42.28 | -79.13 |
|  | WT-4 | -98.02 | -290.78 | 337.94 | -13.19 | -111.21 | 47.16 | -64.05 |
|  | WT-5 | -95.32 | -266.67 | 316.10 | -12.45 | -107.77 | 49.43 | -58.34 |
|  | MT-1 | -106.52 | -213.89 | 261.67 | -14.03 | -120.55 | 47.78 | -72.77 |
|  | MT-2 | -102.07 | -243.14 | 290.14 | -13.41 | -115.48 | 47.00 | -68.48 |
|  | MT-3 | -106.09 | -191.45 | 241.25 | -13.72 | -119.81 | 49.80 | -70.01 |
|  | MT-4 | -106.12 | -258.04 | 305.67 | -14.07 | -120.19 | 47.63 | -72.56 |
|  | MT-5 | -109.66 | -223.46 | 275.33 | -14.48 | -124.14 | 51.87 | -72.27 |

^a)^The unit of the values is kcal/mol; ^b)^WT represents wild-type; ^c)^MT represents mutant.
